# *Colletotrichum higginsianum* effectors exhibit cell to cell hypermobility in plant tissues and modulate intercellular connectivity amongst a variety of cellular processes

**DOI:** 10.1101/2021.01.13.426415

**Authors:** Mina Ohtsu, Joanna Jennings, Matthew Johnston, Xiaokun Liu, Nathan Hughes, Kara Stark, Richard J. Morris, Jeroen de Keijzer, Christine Faulkner

## Abstract

Multicellular organisms exchange information and resources between cells to co-ordinate growth and responses. In plants, plasmodesmata establish cytoplasmic continuity between cells to allow for communication and resource exchange across the cell wall. Some plant pathogens use plasmodesmata as a pathway for both molecular and physical invasion. However, the benefits of molecular invasion (cell-to-cell movement of pathogen effectors) are poorly understood. To begin to investigate this and identify which effectors are cell-to-cell mobile, we performed a live imaging-based screen and identified 15 cell-to-cell mobile effectors of the fungal pathogen *Colletotrichum higginsianum*. Of these, 6 are “hypermobile”, showing cell-to-cell mobility greater than expected for a protein of its size. We further identified 3 effectors that can indirectly modify plasmodesmal aperture. Transcriptional profiling of plants expressing hypermobile effectors implicate them in a variety of processes including senescence, glucosinolate production, cell wall integrity, growth and iron metabolism. However, not all effectors had an independent effect on virulence. This suggests a wide range of benefits to infection gained by the mobility of *C. higginsianum* effectors that likely interact in a complex way during infection.

## Introduction

Cell-to-cell communication is essential for multicellularity and there are a variety of mechanisms by which cells exchange information and resources. Plant cells are surrounded by cell walls but have tunnel-like structures called plasmodesmata which cross the cell wall and directly connect the cytoplasm of adjacent cells to establish the symplast. Small molecules such as sugars, metabolites and hormones can all move between cells through plasmodesmata (Stahl and Simon, 2013; Cheval and Faulkner, 2018; Liu and Chen, 2018), driven by advection and diffusion. Further, larger proteins can pass between cells via mechanisms that involve active components such as protein unfolding and refolding or possible intercellular trafficking motifs/domain (Kragler *et al*., 1998; Taoka *et al*., 2007; Xu *et al*., 2011; Chen *et al*., 2013; Chen *et al*., 2014).

The regulation of plasmodesmata is a critical component of plant-microbe interactions. Many plant immune responses are triggered by cell autonomous recognition of pathogen molecules, but we and others have shown that plasmodesmata dynamically respond to immune signals. We previously found that both fungal and bacterial molecules induce plasmodesmal closure in *Arabidopsis thaliana*. Chitin (from fungal cell walls) and flg22 (from bacterial flagellin) both trigger plasmodesmal closure, regulated by LYSM-CONTAINING GPI-ANCHORED PROTEIN 2 (LYM2) and CALMODULIN-LIKE 41 respectively (Faulkner *et al*., 2013; Xu *et al*., 2017). Observations that plasmodesmal function influences infection outcomes identify that plasmodesmal responses are key to ultimate defence success (Lee *et al*. 2011; Faulkner *et al*., 2013; Caillaud *et al*., 2014; Xu *et al*., 2017). This suggest two possibilities: plasmodesmal regulation is important for the execution of plant immune responses and/or it impairs infection mechanisms deployed by the invading pathogens. The latter implicates pathogen access to the symplast as a critical component of infection.

Plasmodesmata present a route by which microbes can access non-infected cells and tissues. Indeed, there are several examples of pathogens directly using plasmodesmata to facilitate their passage between host cells in a growing infection. The best understood examples of this are viral infections such as *Cucumber mosaic virus* (CMV) and *Tobacco mosaic virus* (TMV) (Heinlein and Epel, 2004). As obligate parasitic pathogens that infect intracellularly, viruses actively target and modify plasmodesmata to translocate their genomes between cells to establish an infection. Interestingly, it has also been revealed that some hemi-biotrophic fungal pathogens, including the rice blast pathogen *Magnaporthe oryza*, pass from cell to cell at plasmodesmal pitfields (Kankanala *et al*., 2007). These demonstrate that plant pathogens across different kingdoms have acquired the capacity to recognize and exploit plasmodesmata as sites of connection between cells to enable the spread of infection.

In order to manipulate host plant systems, plant pathogens secrete an arsenal of proteins, called effectors (Le Fevre *et al*., 2015; Toruño *et al*., 2016). Plasmodesmata allow the spread of soluble molecules, and while effectors can act in the host cell to which they are delivered, soluble effectors also have the potential to move between cells via plasmodesmata. Indeed, *M. oryzae* produces the PWL2 and BAS1 effectors that can move into non-infected cells (Khang *et al*., 2010). This suggests that pathogens exploit plasmodesmata to access and manipulate non-infected cells ahead of the infection front. Further implicating the symplast in infection, the *Fusarium oxysporum* effector Six5 was found to enable cell-to-cell translocation of its co-transcribed effector Avr2 via plasmodesmata (Cao *et al*., 2018). Moreover, both the *Pseudomonas syringae* effector HopO1-1 and the *Phytophthora Brassicaceae* effector RxLR3 target and modify plasmodesmata (Aung *et al*., 2020; Tomcynska *et al*., 2020).

It is not yet fully understood what a microbe gains by accessing the host symplast, or how common cell-to-cell mobility is within effector repertoires. To address these questions, we characterized the cell-to-cell mobility of candidate effectors from the hemi-biotrophic fungal pathogen *Colletotrichum higginsianum*. We used a live imaging-based screen to identify candidate effectors that move cell to cell in plant tissues and identified effectors that are cell-restricted (immobile), move cell to cell to a degree expected for a protein of that size (mobile) and move further than expected (hypermobile). Within the hypermobile effectors, we identified effectors that modify plasmodesmata, consistent with enhanced mobility, and one with a signature of active translocation. Expression of hypermobile effectors in host tissue identified that these three effectors have a differential effect on pathogen virulence and the host transcriptome. The latter identified processes associated with nutrient uptake and defence that illustrate what *C. higginsianum* gains by molecular invasion of host tissues. We conclude that pathogen access to the host symplast is a complex component of the infection strategy of *C. higginsianum* allowing it to manipulate a variety of host processes ahead of the infection front.

## Material and Methods

### Plant material

*Nicotiana benthamiana* plants were grown at 23 °C under long day conditions (16 h: 8 h, light: dark). *Arabidopsis thaliana* were grown on soil at 22 °C under short day conditions (10 h: 14 h, light: dark) or on MS media under short day conditions (10 h: 14 h, light: dark).

### DNA Constructs

Constructs for plant expression were assembled using the Golden Gate cloning method (Engler *et al*., 2008) and all module information is in Table S1. The coding sequences of effector candidates (without predicted signal peptides) were domesticated to remove BpiI and BsaI sites and synthesised as Golden Gate-compatible Level 0 vectors. For subcellular localisation analysis, effector sequences were fused to the N-terminus of GFP and expressed from the CaMV 35S promoter. For the cell-to-cell mobility assay and generating Arabidopsis stable lines, multi-component binary vectors were assembled as outlined in Fig. S1 and Table S1.

### Plant transformation

Transient gene expression in *N. benthamiana* was performed as described (Cheval *et al*., 2020). *Agrobacterium tumefaciens* (GV3101) carrying binary plasmids were infiltrated into *N. benthamiana* leaves at OD_600nm_ = 1.0 × 10^−2^ to check subcellular localisation and at OD_600nm_ = 1.0 × 10^−5^ to generate single cell transformation events for the mobility assay. Samples were imaged 72 h post infiltration. Stable transgenics were made by floral dipping Arabidopsis Col-0.

### Microscopy

Leaf tissue was imaged by confocal microscopy (Zeiss LSM800) with a 20x water-dipping objective (W N-ACHROPLAN 20x/0.5; Zeiss). GFP was excited with a 488 nm solid-state laser and collected at 509-530 nm and dTomato was excited by a 561 nm solid-state laser and collected at 600-640 nm using sequential scanning.

### Image and data analysis

To quantify effector-GFP mobility we recorded the number of fluorescent cells around the transformed cell, identified by NLS-dTomato fluorescence. Data analysis was performed in R statistical computing language v4.0.3 (R Core Team, 2020). The standard curve was generated with data obtained from mobility of GFP, YFPN-GFP, YFPC-GFP and 2xGFP using a quasi-Poisson generalised linear model with a log link function and a Bonferroni corrected p-value < 1× 10^−5^. Effectors were determined to be significantly mobile by the equivalent of a t-test for Poisson distributions (an exact Poisson test) in which the mobility (cell count at 3dpi) is significantly different to the standard curve (*p* <1×10^−5^). For analysis of NLS-dTomato movement in the presence of different effectors we tested the null hypothesis that mobility of NLS-dTomato is 2.5 cells (as observed for the set of standard constructs) and independent of effector mobility using an exact Poisson test.

### Protein extraction and Western blotting

Protein extraction was performed as described (Adachi *et al*., 2019). *N. benthamiana* leaves transiently expressing effectors were collected tissue for protein extraction 3 days post infiltration. Leaf material was homogenised in ice-cold extraction buffer (10% glycerol, 25 mM Tris pH 7.5, 1 mM EDTA, 150 mM NaCl, 2% w/v PVPP, 10 mM DTT, protease inhibitor cocktail (Sigma), 0.2% Igepal (Sigma)) and the soluble fraction was collected by centrifugation at 12,000 x*g* for 10 min. Proteins were separated by SDS-PAGE and transferred PVDF membrane (BioRad). Proteins were detected with 1/5,000-diluted anti-GFP conjugated to HRP (sc-9996; Santa Cruz Biotechnology).

### Microprojectile bombardment

Microprojectile bombardment assays were performed as described (Faulkner *et al*., 2013). Four- to six-week-old expanded leaves of relevant Arabidopsis lines were bombarded with 1 nm gold particles (BioRad) coated with pB7WG2.0-mRFP, using a Biolostic PDS-1000/He particle delivery system (BioRad). Bombardment sites were imaged 24 h post bombardment by confocal microscopy (Zeiss LSM800) with a 10x (EC Plan-NEOFLUAR 10× 0.3; Zeiss) or 20x dry objective (Plan-APOCHROMAT 20x/0.8; Zeiss). Data were collected from at least 2 independent bombardment events, each of which consisted of leaves from at least 3 individual plants. The median normalised mobility (#cells) for different lines was analysed in R statistical computing language v4.0.3 (R Core Team, 2020) by a bootstrap method (Johnston and Faulkner, 2020).

### *C. higginsianum* infection

*C. higginsianum* spores (5 × 10^6^ spores/mL) were drop-inoculated on detached leaves of 4- to 5-week-old Arabidopsis plants on 2% water agar plates. Plates were sealed with parafilm and left for 6 days under short day conditions (10 h: 14 h, light: dark; 25 °C). The area of necrotic lesions was measured in Fiji (Schindelin *et al*., 2012). Lesions were measured for at least 30 plants (2 leaves per plant) and across 3 independent experiments. The mean lesion area for different lines was compared using a bootstrapping method (Johnston and Faulkner, 2020).

### RNAseq analysis and GO term analyses

Leaves 7 and 8 of 4-week-old MS grown Arabidopsis was harvested for RNA extraction. Leaves from 3 plants were pooled for a single replicate and 3 replicates were collected for each genotype. RNA was extracted using RNAeasy Mini Kit (Qiagen) followed by DNase treatment (TurboDNase, ThermoFisher Scientific). Library preparation and Illumina sequencing (PE150, Q30>80%) with 10M paired reads was performed by Novogene. Sequencing reads quality was evaluated using FastQC v0.11.9 (Andrews, 2010) and multiqc v1.9 (Ewels *et al*., 2016). After quality control, trimmomatic v0.39 was used to remove Illumina sequence adapters and low-quality reads. Processed reads were re-assessed with FastQC v0.11.9 and mapped to the Arabidopsis thalina TAIR10 release 37 genome assembly using hisat2 v2.2.0 and samtools v1.11 (Li *et al*., 2009).

Differential expression analysis was performed with DESeq2 v1.20.0 (Love *et al*., 2014) and the R statistical computing language v4.0.3 (R Core Team, 2020). Analysis using the methods described by Love *et al*., 2014. were used to calculate normalised expression values for each gene across all samples. Normalised expression values were compared for all expressed genes in Col-0 to those in the effector expressing lines using a hypergeometric test with the Benjamini and Hochberg False Discovery Rate (FDR) correction. Differentially expressed genes were defined by |log2[fold change]|≥1 and FDR corrected *p*-value < 0.01. Differentially expressed genes were analysed by GoTermFinder (Boyle *et al*., 2004) with a hypergeometric test with a Bonferroni correction to identify biological processes enriched in the samples (adjusted *p*-value>0.01). CirGO (Kuznetsova *et al*., 2019) was used to visualise the results.

## Results

### Identification of candidate cell-to-cell mobile *Colletotrichum higginsianum* effectors

To establish a candidate list of putative *C. higginsianum* effectors (hereafter referred to as effectors) exhibiting cell-to-cell mobility, we mined published transcriptome data covering different infection stages (O’Connell *et al*., 2012; Dallery *et al*., 2017). Many effectors are secreted from pathogens into host plant tissues (Lo Presti *et al*., 2015), and therefore we limited our candidate cell-to-cell mobile effectors as those that encode conventionally secreted proteins with a predicted signal peptide. Gene expression data in O’Connell *et al*. (2012) defines expression profiles during the following stages of growth and infection: *in vitro* appressoria (VA), *in planta* appressoria (PA), biotrophic phase (BP) of infection and necrotrophic phase (NP) of infection. We reasoned that cell-to-cell mobile effectors would be primarily relevant during the penetration (PA) and biotrophic phases (BP) of infection (i.e. when the host tissue is alive) and thus defined candidate cell-to-cell mobile effectors as those that have enhanced expression in PA and BP phases relative to the other 2 stages (i.e. PA/VA, BP/VA, PA/NP, and BP/NP >2). We further limited candidates to those for whichPA reads > 50 and BP reads > 20. These criteria produced a list of 46 candidate effectors.

Plant proteins that are known to be cell-to-cell mobile are typically soluble within the cytoplasm or the nucleus (Kim *et al*., 2002; Gallagher *et al*., 2004; Gallagher and Benfey, 2009; Chen *et al*., 2013) and we made the assumption that *C. higginsianum* cell-to-cell mobile proteins would have similar properties. Thus, to further refine our list of candidate cell-to-cell mobile effectors, we cloned these 46 candidate effectors as fusions to GFP and screened for nucleocytoplasmic and nuclear subcellular localisations by transient transformation of *N. benthamiana*. Of these 46 effector-GFP fusions, 25 showed nucleocytoplasmic localisation but none showed a specific nuclear localisation.

### Live cell screening for cell-to-cell mobility

To assay the cell-to-cell mobility of candidate effectors, we performed a live cell imaging-based screen using transient transformation of single epidermal cells in *N. benthamiana*. For this we used Golden Gate modular cloning (Engler *et al*., 2008) to assemble effector-GFP fusions and a cell transformation marker in a single vector as a dual expression cassette vector (Fig. 1a). For a cell transformation marker, we used dTomato fused to a nuclear localisation sequence (NLS-dTomato), reasoning that the dimerization properties of dTomato would make a protein complex too large to move from cell to cell.

**Fig 1.**
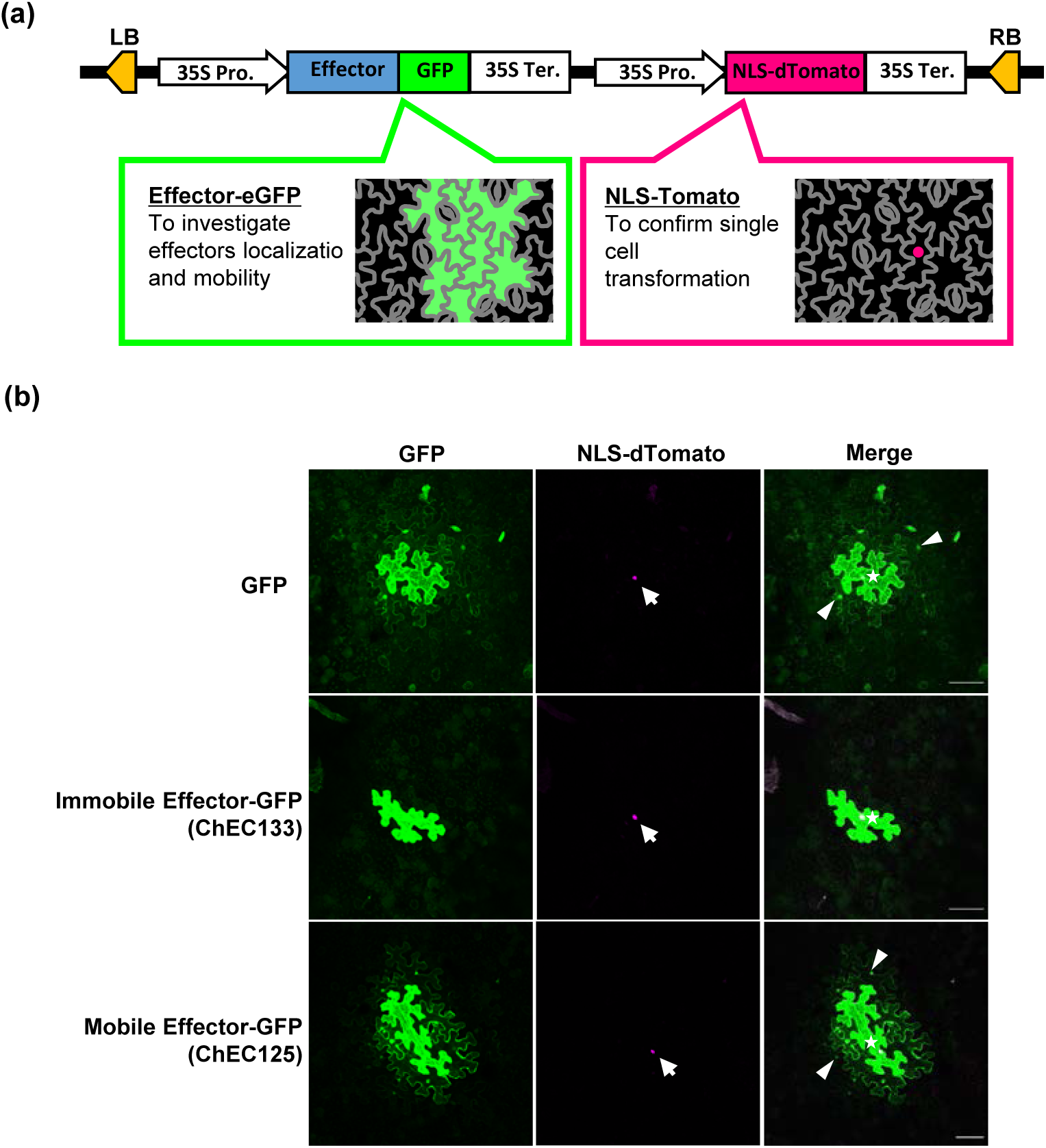
GFP fusions allow detection of cell-to-cell mobility screening of *C. higginsianum* effectors. (a) Schematic of the dual expression module binary vectors used to screen for cell-to-cell mobility. The cartoon represents the patterns of localisation for each module’s gene product expected for a mobile effector-GFP fusion. (b) Projections of confocal z-stacks (8-20 individual focal planes, taken at 5.66 μm intervals) of tissues expressing GFP alone and effector-GFP fusions. Mobility is visible when cells surrounding the transformed cell, marked by NLS-dTomato, show GFP fluorescence. ChEC133-GFP (middle row) shows an example of a cell-restricted, immobile effector while ChEC125-GFP (bottom row) shows an example of mobility to surrounding cells as observed for GFP alone (top panel). Arrows indicate NLS-dTomato fluorescence in the transformed cell, stars represent the transformed cells and arrowheads represent cells the GFP has moved into. Scale bar is 100 µm.

To confirm the utility of NLS-dTomato as a cell transformation marker, we generated constructs that express GFP or 2xGFP with NLS-dTomato (pICH4723.GFP.NLS-dTomato and pICH4723.2xGFP.NLS-dTomato respectively). Agrobacterium infiltration of *N. benthamiana* leaves demonstrated that both fluorophores were expressed in the transformed cell. While GFP was frequently seen to move freely from the transformed cell, both NLS-dTomato and 2xGFP were mostly retained within the transformed cell (Fig. 1 and S1).

Cell-to-cell mobility via plasmodesmata is dependent upon the size of the molecule (Oparka *et al*., 1999; Crawford and Zambryski., 2001). To determine the relationship between size and mobility in *N. benthamiana* leaves in more detail, we assayed the mobility of four proteins of different sizes: GFP (26 kDa), YFPc-GFP (37.2kDa), YFPn-GFP (45 kDa) and 2x GFP (52 kDa) (Fig. S1). We generated single cell transformation sites by low OD600 Agrobacterium infiltration of *N. benthamiana* leaves and counted the number of cells which exhibited GFP surrounding a transformed cell for each construct (marked by NLS-dTomato) (Fig. 1). We fit a quasi-Poisson function through the data which makes minimal assumptions, obtaining a smooth fit through the data that allows for the infrequent mobility of the larger standard proteins (Fig. S1). However, we point out that the data itself does not exclude the existence of a SEL which could be anywhere upwards from YFPc-GFP. This model, with a Bonferroni corrected confidence interval of the mean (*p* < 1×10^−5^), defined a standard curve against which to compare mobility of effectors of varying sizes.

### Cell-to-cell mobility screen identifies mobile and hypermobile effectors

To characterize the cell-to-cell mobility of nucleocytoplasmic effectors, we cloned each of the 25 effectors in a dual expression cassette vector as described (i.e. pICH4723.Effector-GFP.NLS-dTomato) (Fig. 1a). Assaying for cell-to-cell mobility, we observed varying degrees of cell-to-cell mobility for the effectors (i.e. Fig. 1, lower panels); 10 effectors were restricted to the transformed cell (i.e. Fig. 1b, middle panels) in *N. benthamiana* and 15 exhibited cell-to-cell mobility (Table S2).

To determine whether an effector’s mobility was as expected for a soluble protein of that size, we compared effector mobility to the standard curve (Fig. S1b; Fig. S2). An exact Poisson test identified a subset of 7 effectors that had greater than expected mobility, and we defined these as ‘hypermobile’ (Fig. 2). To confirm that this mobility did not arise from cleavage of the effector-GFP fusion (to produce a smaller and thus more mobile protein), we assayed the protein size of the GFP-fused 7 hypermobile effectors expressed in *N. benthamiana* by protein extraction and Western blot analysis. Two strong bands for ChEC130-GFP were observed, suggesting this effector is cleaved in host cells (Fig. S3). However, the other 6 effectors showed little evidence of significant degradation or cleavage (Fig. S3). Thus, we concluded that ChEC123, ChEC124, ChEC125, ChEC128, ChEC129 and ChEC132 are hypermobile.

**Fig 2.**
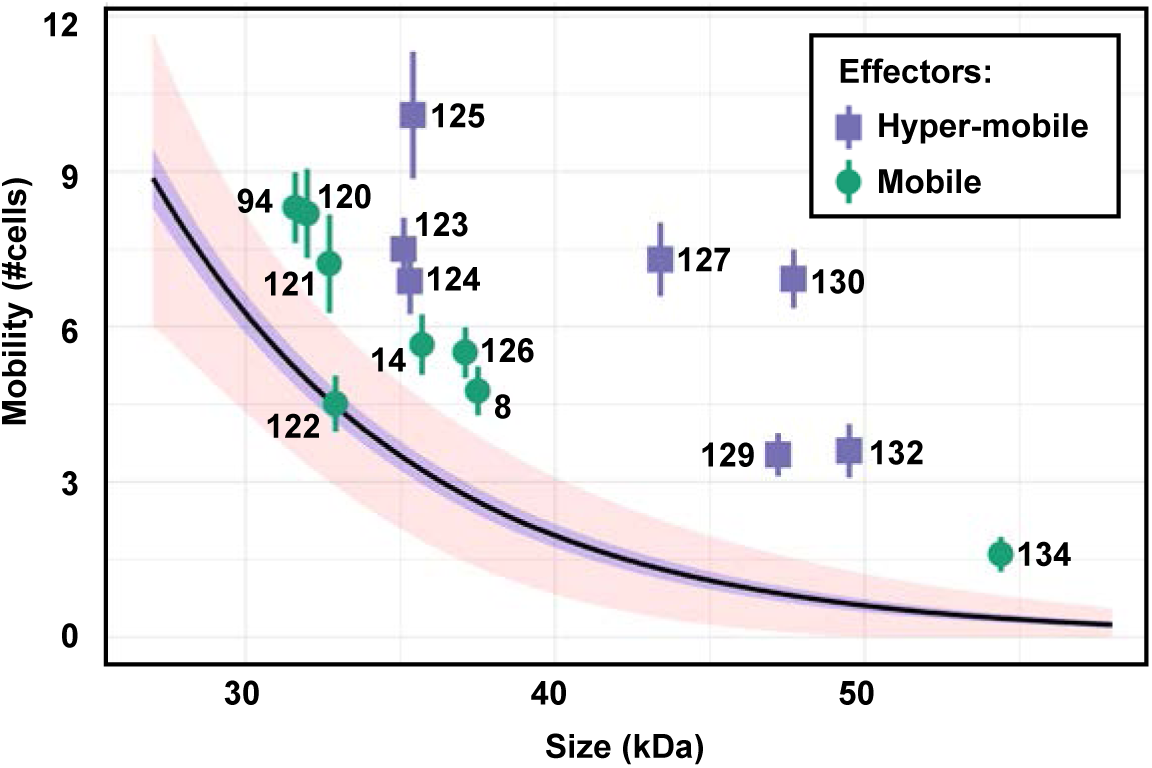
Mobility quantification for nucleocytoplasmic effectors reveals mobile and hypermobile effectors. Mobility (number of cells to which GFP has moved around a transformed cell) of effector-GFP fusions. The standard curve is a quasi-Poisson generalised linear model with a log link function and the Bonferroni corrected confidence interval of the mean (*p* < 1×10^−5^) (red ribbon). Effectors were determined to be significantly mobile (purple squares) by an exact Poisson test indicating the rate of movement is significantly different to the standard curve (*p* < 1×10^−5^). Data are means ± standard error (n >30, *p* < 1 × 10^−5^)

### Mobile effectors can modify plasmodesmal function

The identification of both mobile and hypermobile effectors suggests that *C. higginsianum* might access the host symplast via different mechanisms. Hypermobility suggests three possibilities: that the shapes or Stokes radius of such effectors allows for more efficient translocation through plasmodesmata; that effectors exploit an active translocation mechanism; or effectors modify PD (directly or indirectly) which allows their passage. To address the question of whether mobile and hypermobile effectors modified plasmodesmal function, we exploited the unexpected low-level mobility of NLS-dTomato observed post-collection when the image contrast was adjusted (Fig. S4). NLS-dTomato clearly marked the transformed cell in all cases, but upon increasing the contrast of the images it was observed in an average of 2.5 cells surrounding the brightest transformed cell in size-standard controls (Fig. S4a; Fig. S5). We quantified NLS-dTomato movement when co-expressed with each mobile and hypermobile effector and observed variation in the spread of NLS-dTomato (Fig. 3a; Fig. S4b). To determine if hypermobility is associated with a general increase in mobility that would indicate plasmodesmal regulation, we compared relative mobility (Mob_r_=Mob_observed_/Mob_expected_) to the mobility of NLS-dTomato. An exact Poisson test of this data, using the null hypothesis that mobility of NLS-dTomato is 2.5 cells for all values of Mobr, revealed that ChEC127 and ChEC8 both significantly increase NLS-dTomato mobility (Fig. 3a). This result suggests that ChEC127 and ChEC8 modify plasmodesmal function. Curiously, while ChEC127 is a hypermobile effector, ChEC8 is not, suggesting that despite modifying plasmodesmal function, the effector itself does not have increased translocation. This phenomenon might be explained if ChEC8 binds other proteins in plant cells to increase its effective size.

**Fig 3.**
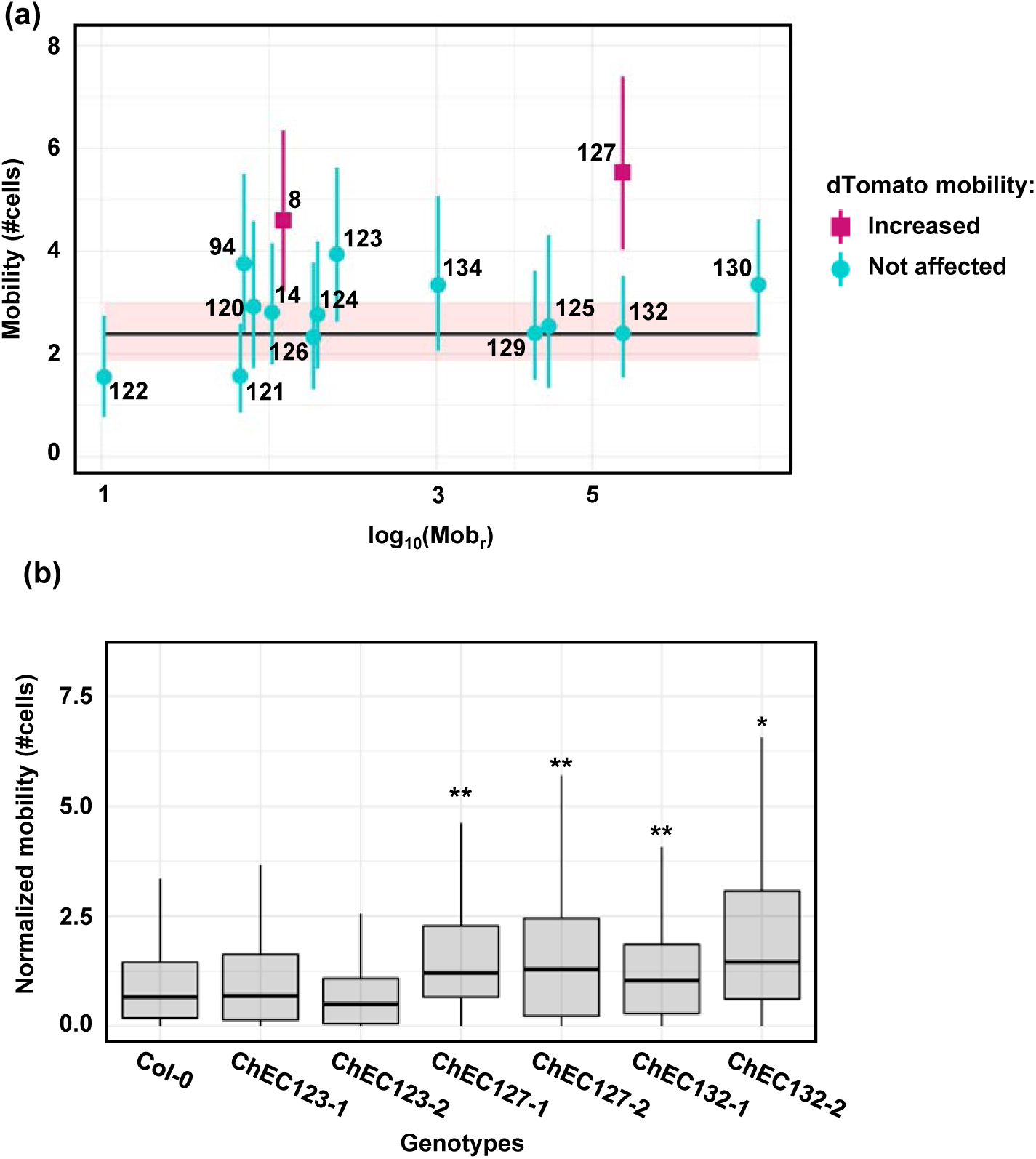
Plasmodesmal regulation by mobile and hypermobile effectors. (a) Mobility of NLS-dTomato plotted against the relative mobility (Mobr) of a co-expressed effector in *N. benthamiana* leaf epidermal cells. The standard curve represents NLS-dTomato mobility in the presence of GFP variants of different sizes with Bonferroni corrected confidence interval of the mean (*p* < 1×10^−5^red ribbon). NLS-dTomato movement was analysed by an exact Poisson test, identifying that ChEC8 and ChEC127 increase NLS-dTomato mobility (b) Mobility of mRFP (number of cells) in Arabidopsis leaf tissue assayed by microprojectile bombardment assays. Independent transgenic lines expressing ChEC127-GFP lines and ChEC132-GFP showed increased mobility of mRFP relative to Col-0. Boxes signifies the upper and lower quartiles, and the whiskers represent the minimum and maximum within 1.5 × interquartile range. The number of bombardment sites (n) counted is ≥ 92. Data was analysed by bootstrap analyses and asterisks indicate statistical significance compared with Col-0 plants (\*\**p* < 0.01 and \**p* < 0.05)

NLS-dTomato is targeted to the nucleus and therefore has a limited pool available in the cytoplasm for cell-to-cell movement. Further, NLS-dTomato is expected to dimerise to form a complex that we expect has reduced mobility as a consequence of increased size. Thus, mobility of this protein is a low sensitivity marker for changes to plasmodesmal function detecting only large changes to plasmodesmal aperture. Therefore, we extended our analysis of plasmodesmal function in the presence of hypermobile mobile effectors to examine the mobility of cytoplasmic mRFP in Arabidopsis, a native host plant of *C. higginsianum*. For this, we generated Arabidopsis lines that stably express fluorescent protein fusions of the hypermobile effectors ChEC123, ChEC127 and ChEC132 (Fig. S6). We performed microprojectile bombardment assays on expanded leaves of two independent lines of each effector (Faulkner *et al*., 2013) and quantified spread of mRFP from transformed cells one day post-bombardment. This data showed that mRFP diffusion in ChEC123-expressing lines was similar to Col-0, while ChEC127 and ChEC132 expressing lines showed increased movement of mRFP relative to the Col-0 control (Fig. 3b). Thus, this data indicates that Ch132 also modifies plasmodesmal function like ChEC127 and ChEC8, and that ChEC123 mediates its translocation by a possible active mechanism that does not involve plasmodesmal modification.

### Heterologous expression of ChEC127 promotes virulence

Pathogen effectors are assumed to positively regulate virulence, facilitating infection success. To assess whether the hypermobile effectors ChEC123, ChEC127 and ChEC132 independently promote virulence, we infected plants constitutively expressing these effectors with *C. higginsianum* and measured the size of disease lesions 6 days post inoculation. These assays showed that both independent transgenic lines that express ChEC127 develop larger disease lesions (Fig. 4), identifying that expression of ChEC127 promotes virulence. Plants expressing ChEC123 or ChEC132 showed no increase in susceptibility (Fig. 4), suggesting that in these conditions neither effector independently promotes virulence.

**Fig 4.**
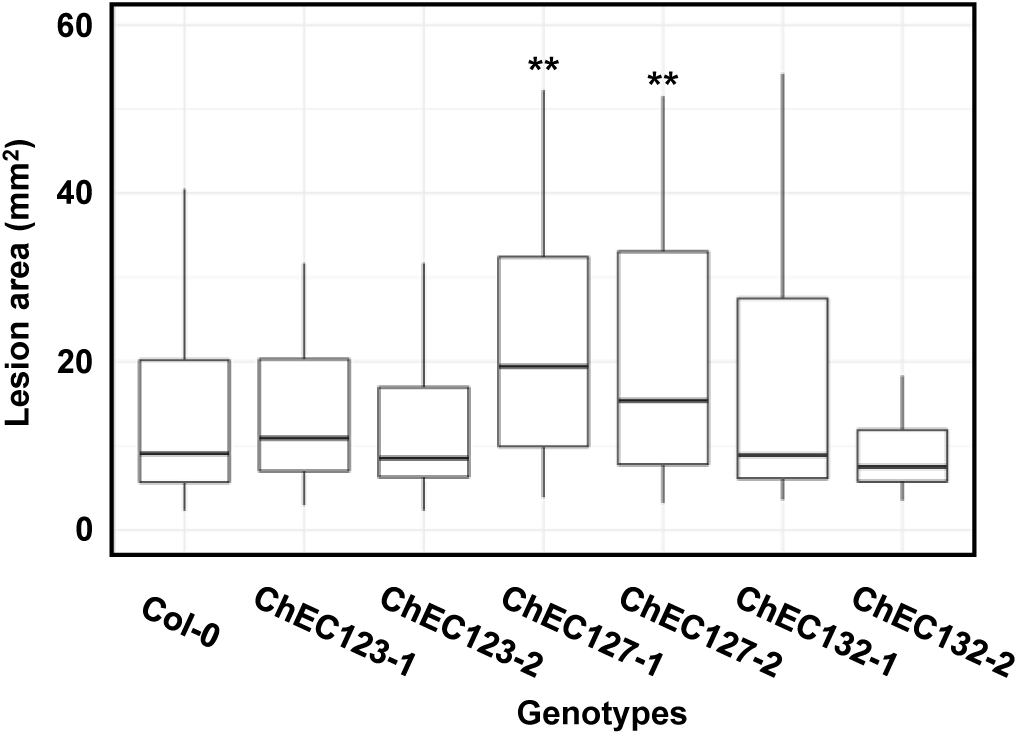
*C. higginsianum* lesion areas are larger in ChEC127 expressing Arabidopsis. Detached leaves from 4-5-week-old Arabidopsis were inoculated with *C. higginisianum* spores and lesion areas were measured 6 dpi (n > 60) and bootstrap analysis of the lesion area means identified that lesions are larger in ChEC127 expressing plants.

### ChEC132 and ChEC123 transcriptionally perturb a variety of host processes

To explore the function of hypermobile effectors, and thus ask what a pathogen might gain from these during infection, we assayed changes to the plant transcriptome induced by ChEC123, ChEC127 and ChEC132. For this, we performed RNAseq analysis of leaf tissue of plants constitutively expressing ChEC123-GFP, ChEC127-GFP and ChEC132-GFP relative to Col-0, defining differentially expressed genes as those for which we detected at least a 2-fold up or down regulation (|log2[fold change]|≥1) and an FDR corrected *p*-value <0.01 To identify processes that are perturbed by these effectors, we used GO Term Finder (Boyle *et al*., 2004) to identify biological processes that are significantly enriched within the differentially expressed genes for each effector.

Despite positively regulating virulence, ChEC127 differentially regulated the expression of only 13 genes (11 up-regulated and 2 down-regulated; Table S4). By contrast, expression of ChEC123 and ChEC132 caused differential expression of 176 and 217 genes respectively (Table S3; S5). GO term analysis indicates ChEC123 up-regulated genes associated with leaf senescence (Fig. 5c) and down regulates genes associated with glycosyl compound catabolism and iron starvation responses and transport (Fig. 5b). Constitutive expression of ChEC132 also induced down regulation of genes associated with iron transport and responses, in addition to cell wall modification, growth, syncytium formation (5 *EXPANSIN* genes also represented in the cell wall loosening GO term), and lipid metabolism (Fig. 5a).

**Fig 5.**
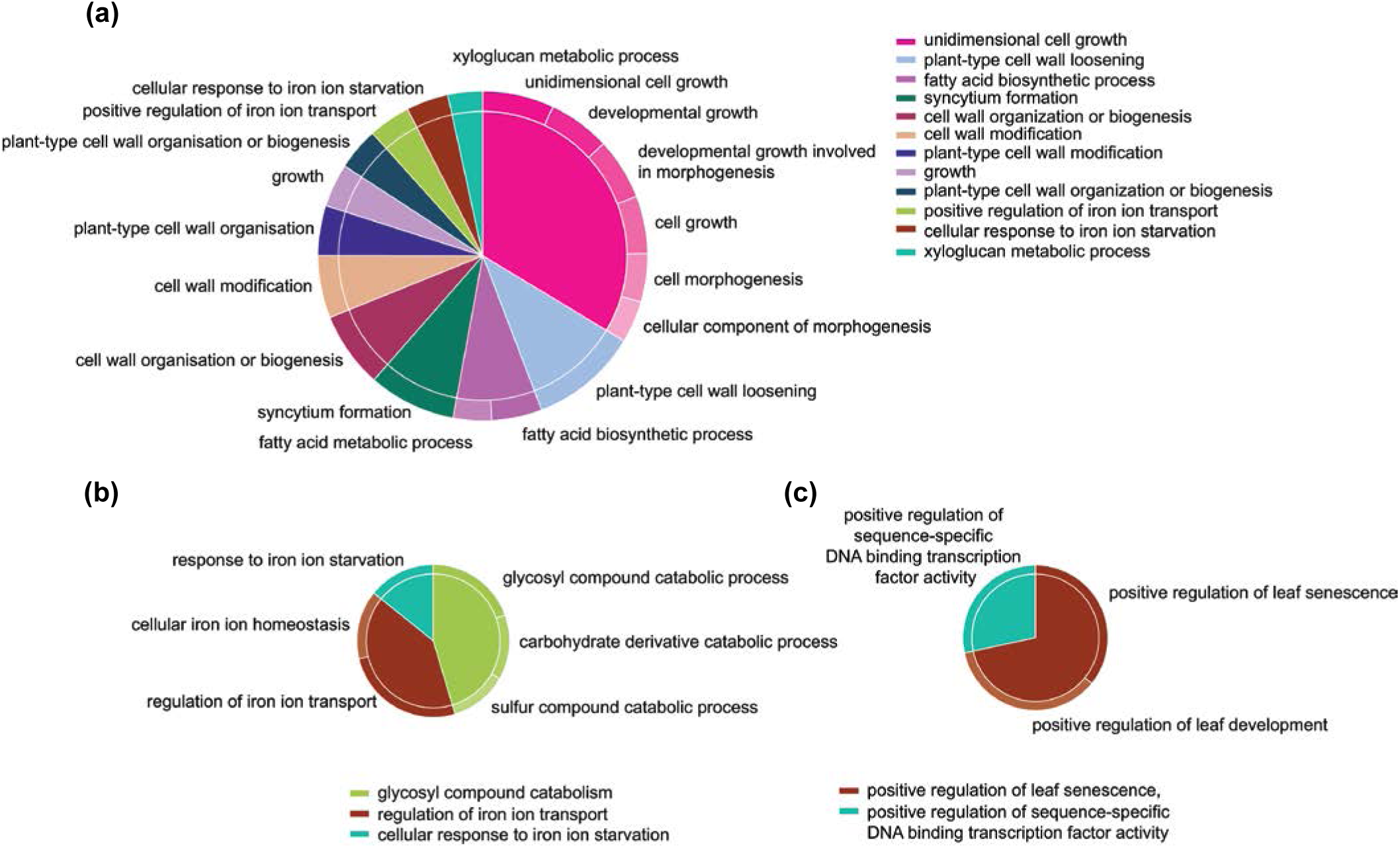
Hypermobile effectors can alter host plant gene expression patterns. CirGO visualisation of GO Terms enriched amongst (a) genes down-regulated by ChEC132, (b) genes down-regulated by ChEC123, and (c) genes upregulated by ChEC123. Slice size represents the proportion of DEGs attributed to this GO Term. The inner ring slices represent ‘parent’ GO terms and the labelled outer ring slices represent ‘child’ GO terms.

## Discussion

Host cell-to-cell connectivity is increasingly identified as a component of plant immunity and pathogen infection. This suggests that pathogen access to non-infected cells is important for the infection strategies of a range of pathogens. Previous studies have identified that specific effectors move cell-to-cell in host tissues (Khang *et al*., 2010; Cao *et al*., 2018), and it was recently suggested that cell-to-cell mobility is a property common to many proteins within an effector repertoire (Li *et al*., 2020). To investigate this further, we performed a screen for cell-to-cell mobility of effectors from the hemi-biotrophic pathogen *C. higginsianum*. We generated a candidate list of secreted effectors from publicly available data and used live-cell imaging to identify a subset of 25 that have a nucleocytoplasmic localisation similar to many known cell-to-cell mobile molecules. To identify mobile effectors, we established a live cell imaging-based screen for cell-to-cell mobility and identified that 60% of nucleocytosolic effectors are cell-to-cell mobile (15/25), with 24% (6/25) showing greater than expected mobility (hyper-mobility) (Fig. 2; Fig. S2; Table S2).

In addition to identifying that cell-to-cell mobility is possible for a range of effectors, our data suggests that effectors from *C. higginsianum* can move through plasmodesmata by different mechanisms. Firstly, we identified a subset of effectors that move ‘passively’ from cell to cell, i.e. they move through plasmodesmata at a rate expected for soluble molecules of the same size (Fig. 2). Like the endogenous plant transcription factor LEAFY (Wu *et al*., 2003), these molecules can be considered to have no mechanism for active translocation and simply move from cell to cell as soluble, freely diffusive molecules. Secondly, we identified hypermobile effectors that modify plasmodesmal function such as ChEC127 and ChEC132. These effectors trigger plasmodesmata opening to a degree such that the effector itself (Fig. 2), as well as other soluble molecules (as observed for mRFP; Fig. 3b), can move faster and further to neighbouring cells. Thirdly, we identified one hypermobile effector, ChEC123, that has no general effect on plasmodesmal function (Fig. 2; Fig. 3). This effector could therefore be considered to have an active and specific mode of translocation to surrounding cells similar to endogenous plant transcription factors such as SHORT-ROOT (SHR) (Nakajima *et al*. 2001), KNOTTED1(KN1) (Lucas *et al*., 1995) or Dof family proteins (Chen *et al*., 2013). This data identifies different mechanisms by which individual effectors can access the symplast, but this data must be considered in the context of infection. Thus, the presence of multiple effectors that modify plasmodesmata during infection raises the possibility that there is a general increase in plasmodesmal aperture that might allow cell-to-cell mobility of effectors that we identified here as immobile.

Our screen revealed that three nucleocytoplasmic effectors can modify plasmodesmal function (ChEC8, ChEC127 and ChEC132), identifying that plasmodesmal regulation might occur during infection via indirect mechanisms. While effectors are expected to positively contribute to virulence, this is not always observable as an independent contribution; effectors may act in concert with other effectors or environmental conditions. In our study, we saw that only ChEC127 significantly and positively contribute to virulence independently as evidenced by increased susceptibility of Arabidopsis plants that express the effector (Fig. 4). Infection is a complex process and interacts with an array of host and environmental variables. Thus, it may not be surprising that excess of a single effector does not promote infection.

To identify host processes that are modulated in a non-cell autonomous way during infection, we performed an RNAseq analysis of plants constitutively expressing the hypermobile effectors ChEC123, ChEC127 and ChEC132 (Table S3-S5). Despite regulating pathogen virulence, ChEC127 induced differential expression of only 13 genes (Table S4), suggesting the mechanism by which it promotes virulence does not involve significant perturbation of gene expression. By contrast, ChEC123 and ChEC132, which did not independently promote virulence, did induce significant changes in gene expression (Table S3; Table S5). ChEC123 downregulated genes associated with glucosinolate production (glycosyl compound catabolism) and iron starvation and transport, indicating it may regulate defence and nutritional processes (Fig. 5b). The same effector up-regulated genes associated with leaf senescence which might contribute to the transition of *C. higginsianum* to the necrotrophic lifestyle (Fig. 5c). ChEC132 down regulated genes associated with iron metabolism (Fig. 5a) but most significantly perturbed processes associated with plant cell wall modification and growth. This raises the possibility that ChEC132 perturbs growth processes, possibly to limit resource consumption by the host. Overall, transcriptional analysis identifies that different host processes can be targeted by cell-to-cell mobile effectors. How manipulation of these processes in cells ahead of the infection front contributes to infection success requires further investigation.

Our screen for cell-to-cell mobility of *C. higginsianum* effectors has identified both mobile and hypermobile effectors. Further, we found evidence that some of these cytoplasmic effectors, both mobile and hypermobile, indirectly regulate plasmodesmata to increase their functional aperture. These observations identify that *C. higginsianum* has complex strategies for accessing the symplast which allows it to perturb processes associated with defence, nutrition and cell structure ahead of hyphal invasion. Evidence of hypermobility in the *C. higginsianum* effector repertoire identifies that exploiting plasmodesmata and cell to cell connectivity to extend pathogen reach is critical for infection.

## Acknowledgements

Research in the Faulkner lab is supported by the European Research Council (725459, “INTERCELLAR”) and the Biotechnology and Biological Research Council (BBS/E/J/000PR9796). MO is supported by the Japan Society for the Promotion of Science, XL was supported by Marie-Curie Fellowship (749755, “HOPESEE”), JJ and NH are supported by the Norwich Research Park Doctoral Training Programme and MJ is supported by the John Innes Foundation.

## Authors Contributions

MO, JJ, XL, KS, and JdK generated materials and performed experiments; MO, JJ, MJ, RJM and NH analysed data; CF managed the project; and MO, JJ and CF wrote the paper with support from all co-authors.

## Data Availability

Raw or normalised cell counts from image analysis is available in Table S1. DESeq2 analysis of RNA sequencing experiments is available in Tables S3-S5.

## Supplementary Figure Legends

**Fig. S1 Mobility of GFP variants of known sizes to generate a standard curve**. Binary vectors with different sized GFP fusions and NLS-Tomato were transiently expressed in 5-week-old *N. benthamiana* and imaged by confocal microscopy 3 dpi: GFP (26 kDa), YFPc-GFP (37.2 kDa), YFPn-GFP (45 kDa) and 2xGFP (52 kDa). (a) Arrows indicate NLS-dTomato fluorescence in the nuclei of transformed cells and arrowheads indicate examples of GFP movements and stars indicate the transformed cell. Each image is a maximum projection of a z-stack comprising 8-20 individual focal planes acquired at 4.61/5.66 μm intervals. Scale bars represent 100 µm. (b) Observed mobility for the various GFP-fusions plotted against their molecular weight. This data was used to define a standard curve with a quasi-Poisson generalised linear model with a log link function. The standard error (purple ribbon) and the Bonferroni corrected confidence interval of the mean (*p* < 1×10^−5^, red ribbon) was calculated. The point density shows the number of replicates at that value.

**Fig. S2 Effector-GFP movement was dependent on effector size**. Raw data of mobility assay showing data spread (summarised in Figure 2) for all effector-GFP fusions. The dot gray level indicates the number of replicates at that value.

**Fig. S3 Stability of hypermobile effector-GFPs in *N. benthamiana* leaves** Five-week-old *N*.*benthamiana* leaves expressing free GFP and effector-GFP fusions were harvested 3dpi. Total proteins were extracted from harvested leaves, separated by SDS-PAGE and were detected using anti GFP antibody. Protein loading was monitored by Coomassie Brilliant Blue (CBB) staining of bands corresponding to the ribulose-1,5-bisphosphate carboxylase large subunit (RBCL).

**Fig. S4 Mobility of NLS-dTomato in effector expressing tissues** (a) Mobility of NLS-dTomato was detected when image display settings were adjusted post-collection. The image on the left shows the imaging data under unsaturated black/white display levels and on the right when the brightness and contrast were enhanced. The image represents a maximum projection of a z-stack comprising 11 individual focal planes. Scale bars represent 50 µm. (b) NLS-dTomato was detected in surrounding cells (arrows) when co-expressed with a variety of effector-GFP fusions. Stars identify the transformed cell. Images are maximum projections of z-stacks comprising 8-20 individual focal planes acquired at an interval of 5.66 μm. Scale bars represent 100 µm.

**Fig. S5 NLS-Tomato moves an average of 2**.**5 cells irrespective of GFP fusion size**. Binary vectors encoding different sized GFP fusions (from 26kDa to 52kDa) and NLS-Tomato were transiently expressed in *N. benthamiana* leaves and imaged by confocal microscopy 3 dpi. The number of cells the NLS-dTomato had moved was counted and a line of best fit generated. The standard error (purple ribbon) and the Bonferroni corrected confidence interval of the mean (*p* < 1×10^−5^) (red ribbon) was calculated for the data. The dot gray level indicates the number of replicates at that value.

**Fig. S6 Expression and localisation of the hypermobile effector in Arabidopsis stable lines** Confocal micrographs of the epidermis of mature leaves of two independent transgenic Arabidopsis lines that express ChEC123, ChEC127 and ChEC132 fused to a fluorescent protein. Each image is a maximum projection of a z-stack comprising 8-20 individual focal planes acquired at a plane interval of 3 μm. Scale bars are 100 µm.

## Supplemental Table legends

**Table S1** Plasmids used and constructed in this study

**Table S2** Nucleocytoplasmic effectors screened for cell-to-cell mobility

**Table S3** DESeq2 analysis of ChEC123-expressing *A. thaliana* compared to Col-0

**Table S4** DESeq2 analysis of ChEC127-expressing *A. thaliana* compared to Col-0

**Table S5** DESeq2 analysis of ChEC132-expressing *A. thaliana* compared to Col-0

**Table S6** Raw data from mobility assays presented in Fig 2, Fig 3 and Fig S2.

## References

Adachi H, Contreras MP, Harant A, Wu CH, Derevnina L, Sakai T, Duggan C, Moratto E, Bozkurt TO, Maqbool A et al. 2019. An N-terminal motif in NLR immune receptors is functionally conserved across distantly related plant species. Elife 8.

Andrews S. 2010. FastQC: a quality control tool for high throughput sequence data. Available online at: http://www.bioinformatics.babraham.ac.uk/projects/fastqc

Aung K, Kim P, Li Z, Joe A, Kvitko B, Alfano JR, He SY. 2020. Pathogenic Bacteria Target Plant Plasmodesmata to Colonize and Invade Surrounding Tissues. Plant Cell 32(3):595–611.

Boyle EI., Weng S., Gollub J., Jin H., Botstein D., Cherry JM., Sherlock G. 2004. GO::TermFinder--open source software for accessing Gene Ontology information and finding significantly enriched Gene Ontology terms associated with a list of genes. Bioinformatics 20:3710–5.

Caillaud MC, Wirthmueller L, Sklenar J, Findlay K, Piquerez SJ, Jones AM, Robatzek S, Jones JD, Faulkner C. 2014. The plasmodesmal protein PDLP1 localises to haustoria-associated membranes during downy mildew infection and regulates callose deposition. PLoS Pathogen 10(10):e1004496–e1004496.

Cao L, Blekemolen MC, Tintor N, Cornelissen BJC, Takken FLW. 2018. The Fusarium oxysporum Avr2-Six5 Effector Pair Alters Plasmodesmatal Exclusion Selectivity to Facilitate Cell-to-Cell Movement of Avr2. Molecular Plant 11(5):691–705.

Chen H, Ahmad M, Rim Y, Lucas WJ, Kim JY. 2013. Evolutionary and molecular analysis of Dof transcription factors identified a conserved motif for intercellular protein trafficking. New Phytol. 198(4):1250–1260.

Chen H, Jackson D, Kim JY. 2014. Identification of evolutionarily conserved amino acid residues in homeodomain of KNOX proteins for intercellular trafficking. Plant Signal Behav. 9(3):e28355–e28355.

Cheval C, Faulkner C. 2018. Plasmodesmal regulation during plant-pathogen interactions. New Phytol. 217(1):62–67.

Cheval C, Samwald S, Johnston MG, de Keijzer J, Breakspear A, Liu X, Bellandi A, Kadota Y, Zipfel C, Faulkner C. 2020. Chitin perception in plasmodesmata characterizes submembrane immune-signaling specificity in plants. Proc. Natl. Acad. Sci. U. S. A. 117(17):9621–9629.

Crawford K, Zambryski P. 2001. Non-targeted and targeted protein movement through plasmodesmata in leaves in different developmental and physiological states. Plant Physiol. 125(4):1802–1812.

Dallery JF, Lapalu N, Zampounis A, Pigné S, Luyten I, Amselem J, Wittenberg AHJ, Zhou S, de Queiroz MV, Robin GP et al. 2017. Gapless genome assembly of Colletotrichum higginsianum reveals chromosome structure and association of transposable elements with secondary metabolite gene clusters. BMC Genomics 18(1):667–667.

Engler C., Kandzia R., Marillonnet S. 2008. A one pot, one step, precision cloning method with high throughput capability. PLoS One 3(11):e3647.

Ewels P., Magnusson M., Lundin S., Käller M. 2016. MultiQC: summarize analysis results for multiple tools and samples in a single report. Bioinformatics 32(19):p 3047–3048.

Farmer E., Mousavi S., Lenglet A. 2013. Leaf numbering for experiments on long distance signalling in Arabidopsis protocol exchange doi:10.1038/protex.2013.071

Faulkner C, Petutschnig E, Benitez-Alfonso Y, Beck M, Robatzek S, Lipka V, Maule AJ. 2013. LYM2-dependent chitin perception limits molecular flux via plasmodesmata. Proc. Natl. Acad. Sci. U. S. A. 110(22):9166–9170.

Gallagher KL, Benfey PN. 2009. Both the conserved GRAS domain and nuclear localization are required for SHORT-ROOT movement. Plant J. 57(5):785–797.

Gallagher KL, Paquette AJ, Nakajima K, Benfey PN. 2004. Mechanisms regulating SHORT-ROOT intercellular movement. Curr. Biol. 14(20):1847–1851.

Heinlein M, Epel BL. 2004. Macromolecular transport and signaling through plasmodesmata. Int. Rev. Cytol. 235:93–164.

Kankanala P, Czymmek K, Valent B. 2007. Roles for rice membrane dynamics and plasmodesmata during biotrophic invasion by the blast fungus. Plant Cell 19(2):706–724.

Johnston MG, Faulkner C. 2020. A bootstrap approach is a superior statistical method for the comparison of cell-to-cell movement data. New Phytol. doi: 10.1111/nph.17159.

Khang CH, Berruyer R, Giraldo MC, Kankanala P, Park SY, Czymmek K, Kang S, Valent B. 2010. Translocation of Magnaporthe oryzae effectors into rice cells and their subsequent cell-to-cell movement. Plant Cell 22(4):1388–1403.

Kim JY, Yuan Z, Cilia M, Khalfan-Jagani Z, Jackson D. 2002. Intercellular trafficking of a KNOTTED1 green fluorescent protein fusion in the leaf and shoot meristem of Arabidopsis. Proc. Natl. Acad. Sci. U. S. A 99(6):4103–4108.

Kragler F, Monzer J, Shash K, Xoconostle-Cázares B, Lucas WJ. 1998. Cell-to-cell transport of proteins: requirement for unfolding and characterization of binding to a putative plasmodesmal receptor. Plant J. 15(3):367–381.

Le Fevre R, Evangelisti E, Rey T, Schornack S. 2015. Modulation of Host Cell Biology by Plant Pathogenic Microbes. Annu. Rev. of Cell and Dev. Biol. 31(1):201–229.

Lee JY, Wang X, Cui W, Sager R, Modla S, Czymmek K, Zybaliov B, van Wijk K, Zhang C, Lu H et al. 2011. A plasmodesmata-localized protein mediates crosstalk between cell-to-cell communication and innate immunity in Arabidopsis. Plant Cell 23(9):3353–3373.

Li H., Handsaker B., Wysoker A., Fennell T., Ruan J., Homer N., Marth G., Abecasis G. Durbin R., 1000 Genome Project Data Processing Subgroup. 2009. The Sequence alignment/map (SAM) format and SAMtools. Bioinformatics 25:2078–9.

Li Z, Variz H, Chen Y, Liu S, Aung K. 2020. Plasmodesmata-dependent intercellular movement of bacterial effectors. bioRxiv doi:10.1101/2020.12.10.420240

Liu L, Chen X. 2018. Intercellular and systemic trafficking of RNAs in plants. Nat. Plants 4(11):869–878.

Love MI., Huber W., Anders S. 2014. Moderated estimation of fold change and dispersion for RNA-seq data with DESeq2. Genome Biology 15:550.

Lucas W, Bouché-Pillon S, Jackson D, Nguyen L, Baker L, Ding B, Hake S. 1995. Selective trafficking of KNOTTED1 homeodomain protein and its mRNA through plasmodesmata. Science 270(5244):1980–1983.

Nakajima K, Sena G, Nawy T, Benfey PN. 2001. Intercellular movement of the putative transcription factor SHR in root patterning. Nature 413(6853):307–311.

O’Connell RJ, Thon MR, Hacquard S, Amyotte SG, Kleemann J, Torres MF, Damm U, Buiate EA, Epstein L, Alkan N et al. 2012. Lifestyle transitions in plant pathogenic Colletotrichum fungi deciphered by genome and transcriptome analyses. Nat. Genet. 44(9):1060–1065.

Oparka KJ, Roberts AG, Boevink P, Cruz SS, Roberts I, Pradel KS, Imlau A, Kotlizky G, Sauer N, Epel B. 1999. Simple, but Not Branched, Plasmodesmata Allow the Nonspecific Trafficking of Proteins in Developing Tobacco Leaves. Cell 97(6):743–754.

Presti LL., Lanver D., Schweizer G., Tanaka S., Liang L., Tollot M., Zuccaro A., Reissmann S., Kahmann R. 2015. Fungal effectors and plant susceptibility. Annu. Rev. Plant Biol. 66:513–45.

R Core Team. 2020. R: a language and environment for statistical computing, v4.0.3. R Foundation for Statistical Computing, Vienna. https://www.R-project.org

Schindelin J, Arganda-Carreras I, Frise E, Kaynig V, Longair M, Pietzsch T, Preibisch S, Rueden C, Saalfeld S, Schmid B,. et al. 2012. Fiji: an open-source platform for biological-image analysis. Nat. Methods. 9 (7):676–82.

Schlicker A., Domingues FS., Rahnenführer J., Lengauer T. 2006. A new measure for functional similarity of gene products based on Gene Ontology. BMC Bioinformatics 7:302.

Stahl Y, Simon R. 2013. Gated communities: apoplastic and symplastic signals converge at plasmodesmata to control cell fates. J. Exp. Bot. 64(17):5237–5241.

Supek F, Bošnjak M, Škunca N, Šmuc T. 2011. REVIGO summarizes and visualizes long lists of Gene Ontology terms. PLoS ONE 6(7):e21800.

Taoka K, Ham BK, Xoconostle-Cázares B, Rojas MR, Lucas WJ. 2007. Reciprocal phosphorylation and glycosylation recognition motifs control NCAPP1 interaction with pumpkin phloem proteins and their cell-to-cell movement. Plant Cell 19(6):1866–1884.

Tomczynska I, Stumpe M, Doan TG, Mauch F. 2020. A Phytophthora effector protein promotes symplastic cell-to-cell trafficking by physical interaction with plasmodesmata-localised callose synthases. New Phytol. 227(5):1467–1478.

Toruño TY, Stergiopoulos I, Coaker G. 2016. Plant-Pathogen Effectors: Cellular Probes Interfering with Plant Defenses in Spatial and Temporal Manners. Annu. Rev. of Phytopathol. 54(1):419–441.

Wu X, Dinneny JR, Crawford KM, Rhee Y, Citovsky V, Zambryski PC, Weigel D. 2003. Modes of intercellular transcription factor movement in the Arabidopsis apex. Development 130(16):3735–3745.

Xu B, Cheval C, Laohavisit A, Hocking B, Chiasson D, Olsson TSG, Shirasu K, Faulkner C, Gilliham M. 2017. A calmodulin-like protein regulates plasmodesmal closure during bacterial immune responses. New Phytol. 215(1):77–84.

Xu X, Wang J, Xuan Z, Goldshmidt A, Borrill P, Hariharan N, Kim J, Jackson D. 2011. Chaperonins facilitate KNOTTED1 cell-to-cell trafficking and stem cell function. Science 333(6046):1141–1144

