## Supplementary Figures for "*Colletotrichum higginsianum* effectors exhibit cell to cell hypermobility in plant tissues and modulate intercellular connectivity amongst a variety of cellular processes"

### Fig S1

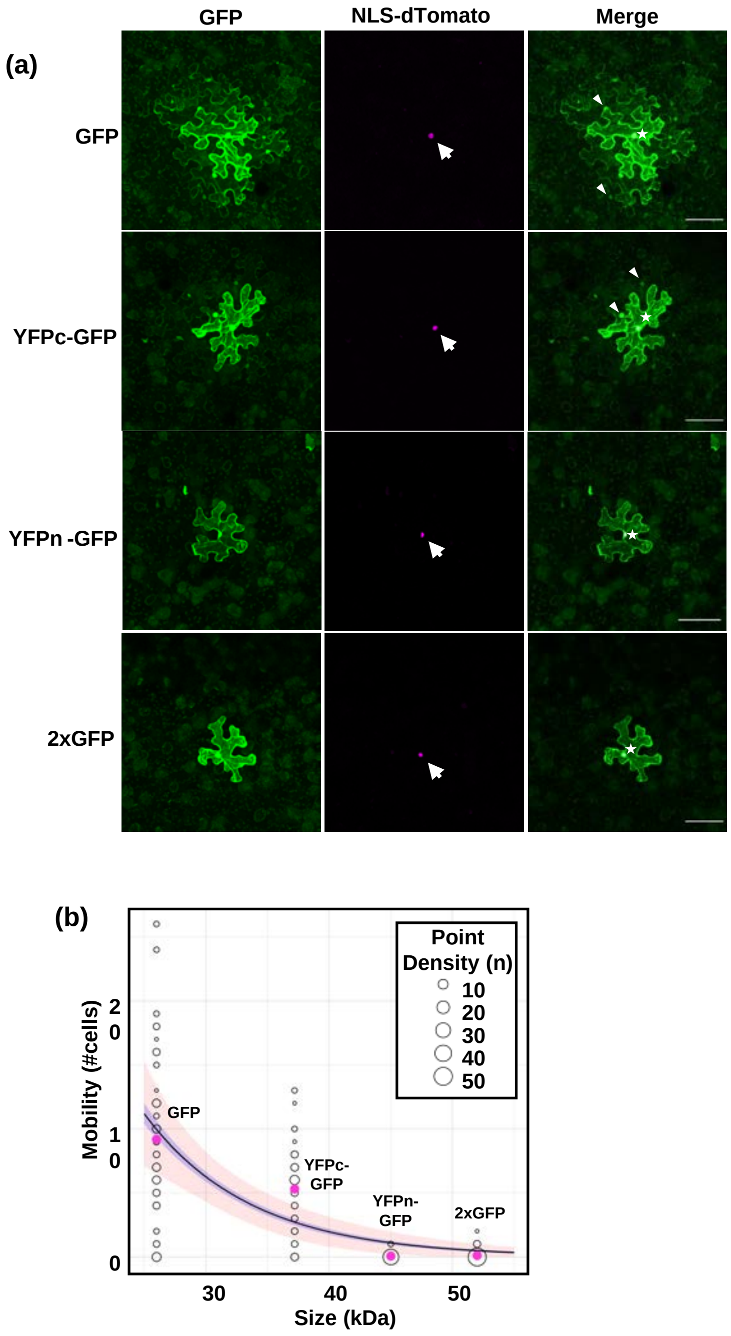


**Fig. S1 Mobility of GFP variants of known sizes to generate a standard curve.** Binary vectors with different sized GFP fusions and NLS-Tomato were transiently expressed in 5-week-old *N. benthamiana* and imaged by confocal microscopy 3 dpi: GFP (26 kDa), YFPc-GFP (37.2 kDa), YFPn-GFP (45 kDa) and 2xGFP (52 kDa). (a) Arrows indicate NLS-dTomato fluorescence in the nuclei of transformed cells and arrowheads indicate examples of GFP movements and stars indicate the transformed cell. Each image is a maximum projection of a z-stack comprising 8-20 individual focal planes acquired at 4.61/5.66 µm intervals. Scale bars represent 100 µm. (b) Observed mobility for the various GFP-fusions plotted against their molecular weight. This data was used to define a standard curve with a quasi-Poisson generalised linear model with a log link function. The standard error (purple ribbon) and the Bonferroni corrected confidence interval of the mean ($p < 1x10^{-5}$, red ribbon) was calculated. The point density shows the number of replicates at that value.

### Fig S2

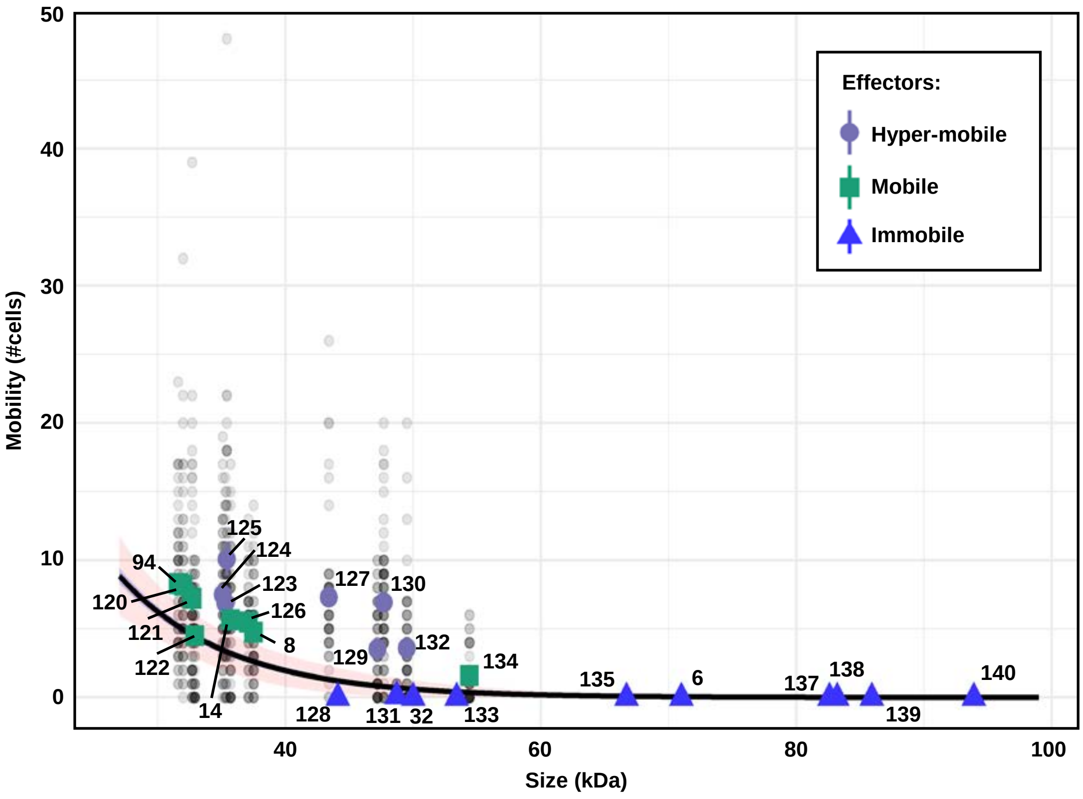


**Fig. S2 Effector-GFP movement was dependent on effector size.** Raw data of mobility assay showing data spread (summarised in Figure 2) for all effector-GFP fusions. The dot gray level indicates the number of replicates at that value.

Fig S3

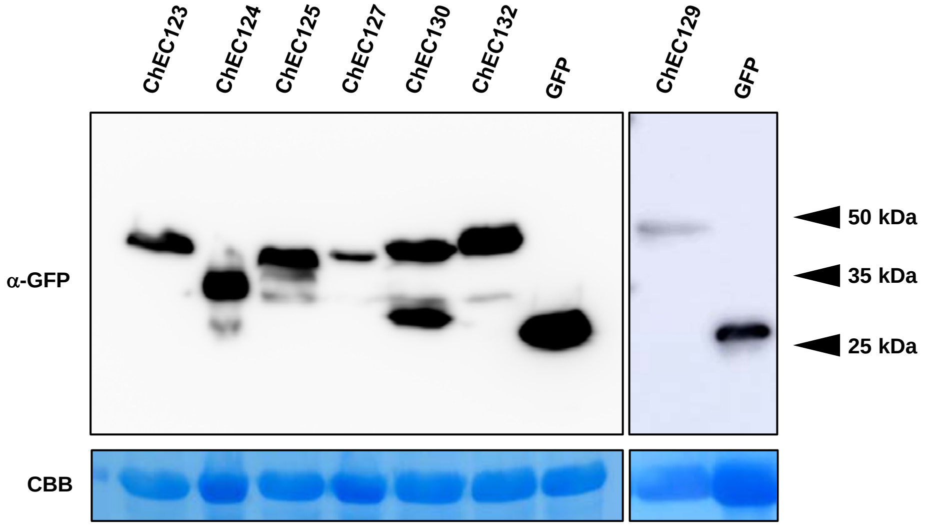


**Fig. S3 Stability of hypermobile effector-GFPs in *N. benthamiana* leaves** Five-week-old *N.benthamiana* leaves expressing free GFP and effector-GFP fusions were harvested 3dpi. Total proteins were extracted from harvested leaves, separated by SDS-PAGE and were detected using anti GFP antibody. Protein loading was monitored by Coomassie Brilliant Blue (CBB) staining of bands corresponding to the ribulose-1,5-bisphosphate carboxylase large subunit (RBCL).

.

Fig S4

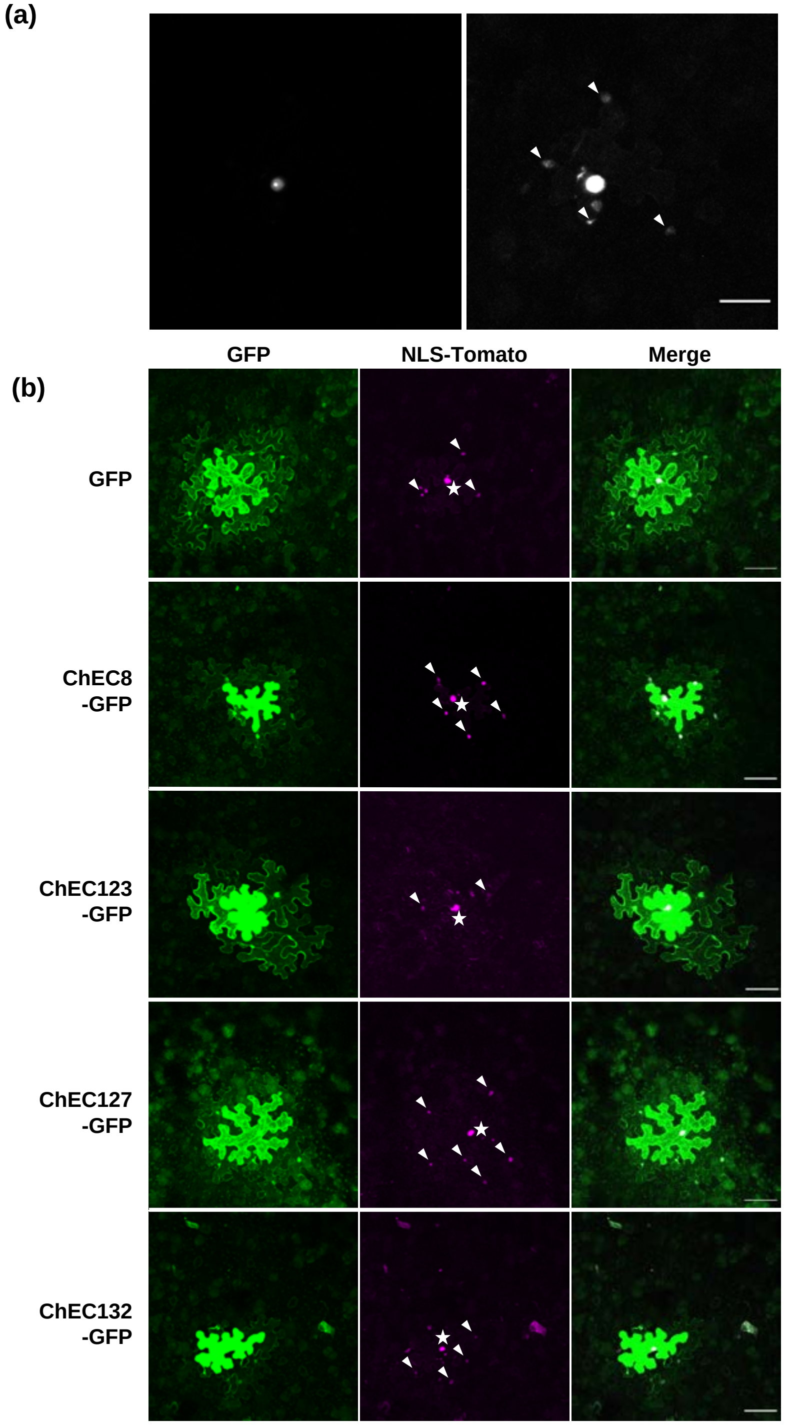


**Fig. S4 Mobility of NLS-dTomato in effector expressing tissues** (a) Mobility of NLS-dTomato was detected when image display settings were adjusted post-collection. The image on the left shows the imaging data under unsaturated black/white display levels and on the right when the brightness and contrast were enhanced. The image represents a maximum projection of a z-stack comprising 11 individual focal planes. Scale bars represent 50 µm. (b) NLS-dTomato was detected in surrounding cells (arrows) when co-expressed with a variety of effector-GFP fusions. Stars identify the transformed cell. Images are maximum projections of z-stacks comprising 8-20 individual focal planes acquired at an interval of 5.66 µm. Scale bars represent 100 µm.

### Fig S5

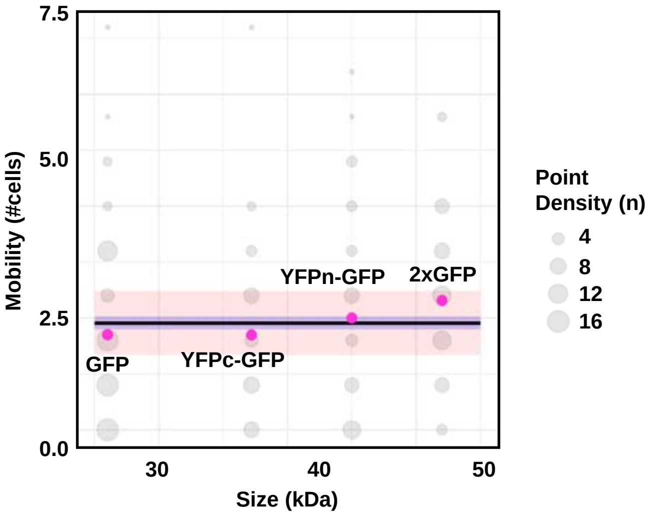


**Fig. S5 NLS-Tomato moves an average of 2.5 cells irrespective of GFP fusion size.** Binary vectors encoding different sized GFP fusions (from 26kDa to 52kDa) and NLS-Tomato were transiently expressed in *N. benthamiana* leaves and imaged by confocal microscopy 3 dpi. The number of cells the NLS-dTomato had moved was counted and a line of best fit generated. The standard error (purple ribbon) and the Bonferroni corrected confidence interval of the mean ($p < 1x10^{-5}$) (red ribbon) was calculated for the data. The dot gray level indicates the number of replicates at that value.

Fig S6

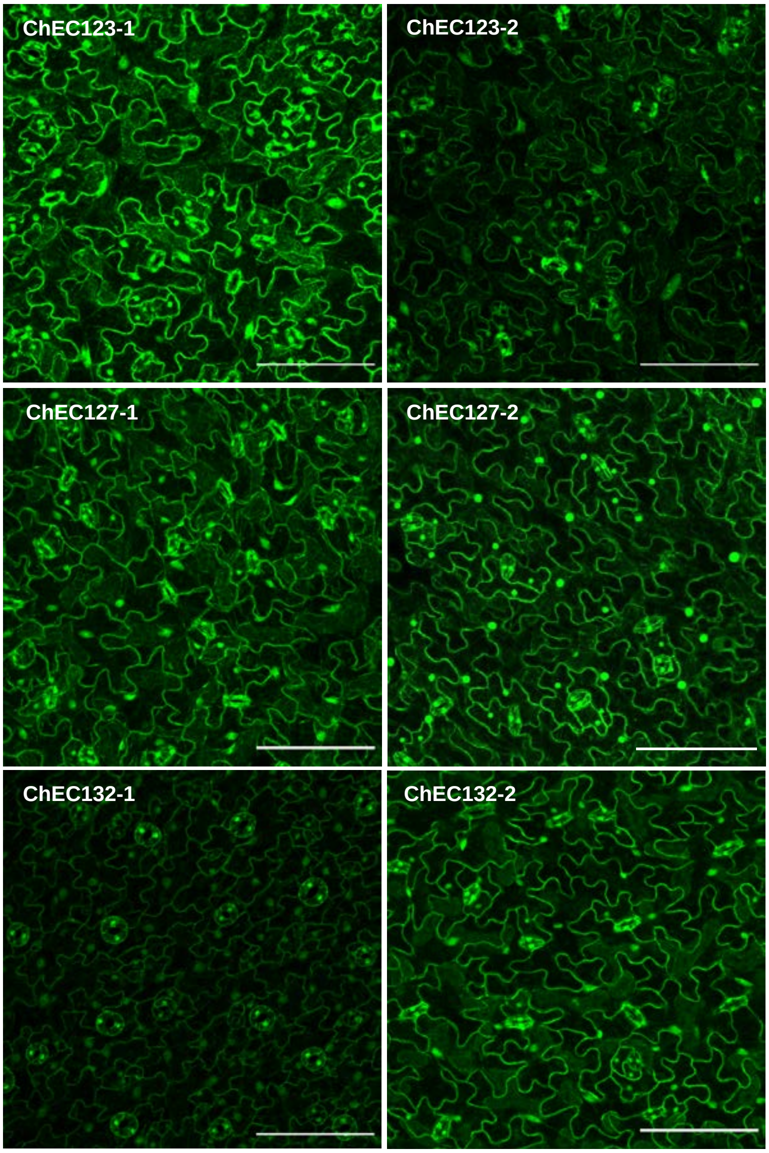


**Fig. S6 Expression and localisation of the hypermobile effector in Arabidopsis stable lines** Confocal micrographs of the epidermis of mature leaves of two independent transgenic Arabidopsis lines that express ChEC123, ChEC127 and ChEC132 fused to a fluorescent protein. Each image is a maximum projection of a z-stack comprising 8-20 individual focal planes acquired at a plane interval of 3 µm. Scale bars are 100 µm.
